# Dynamics of neural microstates in the VTA-striatal-prefrontal loop during novelty exploration in the rat

**DOI:** 10.1101/2020.08.27.270249

**Authors:** A. Mishra, N. Marzban, M. X Cohen, B. Englitz

## Abstract

EEG microstates refer to quasi-stable spatial patterns of scalp potentials, and their dynamics have been linked to cognitive and behavioral states. Neural activity at single and multiunit levels also exhibit spatiotemporal coordination, but this spatial scale is difficult to relate to EEG. Here, we translated EEG microstate analysis to triple-area local field potential (LFP) recordings from up to 192 electrodes in rats to investigate the mesoscopic dynamics of neural microstates within and across brain regions.

We performed simultaneous recordings from the prefrontal cortex (PFC), striatum (STR), and ventral tegmental area (VTA) during awake behavior (object novelty and exploration). We found that the LFP data can be accounted for by multiple, recurring, quasi-stable spatial activity patterns with an average period of stability of ~60-100 ms. The top four maps accounted for 60-80% of the total variance, compared to ~25% for shuffled data. Cross-correlation of the microstate time-series across brain regions revealed rhythmic patterns of microstate activations, which we interpret as a novel indicator of inter-regional, mesoscale synchronization. Furthermore, microstate features, and patterns of temporal correlations across microstates, were modulated by behavioural states such as movement and novel object exploration. These results support the existence of a functional mesoscopic organization across multiple brain areas, and open up the opportunity to investigate their relation to EEG microstates, of particular interest to the human research community.

**Significance Statement:** The coordination of neural activity across the entire brain has remained elusive. Here we combine large-scale neural recordings at fine spatial resolution with the analysis of microstates, i.e. short-lived, recurring spatial patterns of neural activity. We demonstrate that the local activity in different brain areas can be accounted for by only a few microstates per region. These microstates exhibited temporal dynamics that were correlated across regions in rhythmic patterns. We demonstrate that these microstates are linked to behavior and exhibit different properties in the frequency domain during different behavioural states. In summary, LFP microstates provide an insightful approach to studying both mesoscopic and large-scale brain activation within and across regions.

## Introduction

The coordination of neural activity across the entire brain has remained incompletely understood. At the same time, recent research has demonstrated the far-reaching influence of behavior and other state changes on the activity in distant and substantial parts of the brain (McGinley et al., 2015; Musall et al., 2019) Previous work has demonstrated that neural activity is structured into short-lived epoques of coordinated activity on the *local scale*, e.g., of small ensembles of cells (Luczak et al., 2009, 2015) and on the *large scale*, e.g., measurable with non-invasive techniques (Lehmann et al., 1987a; Pascual-Marqui et al., 1995).

On the local level, transient spatiotemporal patterns have been observed in visual cortex (Chiu and Weliky, 2001; Kenet et al., 2003; Fiser et al., 2004; Arieli et al., 1995; Omer et al., 2019), auditory cortex (Luczak et al., 2009; Sakata and Harris, 2009) and in other cortical areas in the awake, anesthetized and sleep state in various species using different recording techniques (Mao et al., 2001; Petersen et al., 2003; Battaglia et al., 2004; Massimini et al., 2004).

On the large-scale level, electroencephalography (EEG) has been a valuable tool to study whole-brain activity in humans. Here, transient spatiotemporal patterns have also been identified and termed EEG microstates. They are used to study human cognition, for example visual processing (Britz and Michel, 2011), perceptual awareness (Britz et al., 2014), neuropsychiatric disorders including schizophrenia (Dierks et al., 1997; Lehmann et al., 2005; Kindler et al., 2011), and resting-state networks (Musso et al., 2010; Yuan et al., 2012; Khanna et al., 2015).

The relation between large-scale EEG microstates and small-scale neural population spike correlations is incompletely understood, in part because different analysis methods are applied at different scales (Alishbayli et al., 2019). In the present study, we combined multichannel recordings (up to 192 electrodes across three brain areas) with the analysis techniques developed in the context of EEG microstates, to investigate the dynamics of neural activity on the meso-scale, i.e. in local field potentials (LFPs). We recorded LFPs simultaneously from the prefrontal cortex (PFC), striatum (STR) and ventral tegmental area (VTA) in rats, while they moved freely in an arena in which novel objects were introduced.

We observed reliable LFP microstates in each region that shared characteristics previously identified in EEG microstates, including robustness, temporal dynamics, temporal continuity, and relation to global field power (Khanna et al., 2015; Milz et al., 2016; Mishra et al., 2020). Using concepts from EEG microstate analysis, we verified the validity of these meso-scale, LFP microstates. For example, the LFP microstates explain a large proportion of the variance of the signal, in contrast to their temporally shuffled surrogates. Furthermore, we linked different microstate properties like *Occurrence rate, Temporal coverage*, and *State duration* to behavioural states. A striking finding was that LFP microstates appear to be coordinated across regions: microstate time-series from distant brain regions exhibited distinct cross-correlation patterns that depended on the behavioural state, suggesting functional interactions of these brain regions, manifested by the microstate activation patterns. Functional coupling of these regions is a well established result (Gao et al., 2007; Zhang et al., 2016), which can be analyzed from a larger perspective using LFP microstates. Thus we propose that the EEG microstate method is an insightful approach to identify, define and characterize more local, transient states of neural activation. As proposed by (Luczak et al., 2015), these microstates might form a vocabulary of neural activation that could help structure our understanding of whole brain dynamics.

## Materials and Methods

All experimental procedures were performed in accordance with the EU directive on animal experimentation (2010/63/**EU**), and the Dutch nationally approved ethics project 2015-0129. All recordings were performed in the lab of MXC. We included four male Long-Evans TH:Cre rats (~3 months old, weight: 350-450 g at time of recordings). Rats were housed singly in Makrolon type III cages (40 × 30 × 35 cm; UNO B.V., Zevenaar, The Netherlands) in a room with controlled temperature and humidity (21 ± 2°C, 60 ± 15%). Animals were kept on 12 hour light/dark cycles, and the experiments were performed under the light phase.

### Electrode implants

Each silicon probe was loaded on a plastic 3D-printed drive purchased from 3Dneuro (https://www.3dneuro.com). The 3Drives were lightweight (0.43 g), with a small footprint and base dimensions of 5 × 6 mm, height: 16.7 mm allowing the travel distance of 9 mm and resolution of 62.5 μm per 1/4 turn. Rats were anesthetized with 5% isoflurane (2-3% maintenance) and placed in a stereotaxic frame. They were given a dose of Rimadyl/Carprofen (0.5 mg/kg) and Xylocaine (maximum 0.4 ml) subcutaneously at the incision site. Small craniotomies on the skull were made above the VTA, STR, and PFC. The animals were implanted with custom designed high-density silicon probes (NeuroNexus, Ann Arbor, USA) with specific electrode layout of densely packed recording sites designed for each target region. In PFC and STR, each probe contained four laminar shanks, having 16 recording electrodes per shank in forms of tetrodes for spike recordings and single sites for LFP recordings, thus creating 64 recording sites per probe. 64 electrodes covered an area of 1 x 2 mm with typical spacing of 225um in each shank and 330 um between shanks in PFC. STR electrodes also covered an area of 1 x 2 mm with the same shank distance (330 um). However, two shanks contained only tetrodes and two shanks had only single sites with typical spacing of 130um between single sites and 660um between tetrodes. VTA contained 8 shanks of 8 electrodes each and covered an area of 1.5 x 0.14 mm. All electrodes were referenced to a reference screw implanted on the skull over the cerebellum. Implants were surrounded by and grounded to a copper mesh faraday cage built around the implants and was fixed to the skull surface using dental cement (Vandecasteele et al., 2012). Rats were monitored daily for one week following surgery and recordings began at least one week after surgery. After recovery (minimum 7 days), the electrodes were advanced into the brain toward a final DV of 4 mm for PFC and 5 mm for striatum, and 8 mm for the VTA.

### Experiment design and apparatus

Animals were handled and trained for one week to freely move and get habitualized in a black plastic square box (60×40×40 cm) open field covered with beddings in the bottom of the enclosure. The square enclosure was placed on an electrically grounded faraday cage inside the recording room. In each session, rats were allowed to move freely in the box. A novel object (e.g., a cup or a toy) was presented in the middle of the box in the second session and the same object was repeated in the fourth session (Figure 4A). Rats were allowed to explore the novel object freely. A camera was placed above the box to track movement. Camera resolution was 320×240 (width x height) and captured at a framerate of 30 Hz. The alignment between camera and OpenEphys acquisition system was implemented using event markers. All sessions within a trial were recorded without any inter-session delay. Maximum of one trial per animal was recorded on a single day. There were 27 recording trials in total.

### Recording and cleaning of electrophysiology data

Electrophysiological data were sampled at 30 kHz via OpenEphys hardware and software solutions (Siegle et al., 2017). Offline, data were downsampled to 1 kHz and stored in the EEGLAB (Delorme and Makeig, 2004) format in MATLAB (The Mathworks, Natick, USA). Excessively noisy data segments were removed manually, and excessively channels were removed from the data based on visual inspection of raw signals and variance thresholds (6.6 ± 5.2 channels per dataset per region). Thereafter, average referencing was performed separately for the set of electrodes located in each region. The datasets were then bandpass filtered in the range of 1-30 Hz, using a Hamming windowed sinc FIR filter, to be consistent with the common practice in EEG microstate analysis (Milz et al., 2017).

### Microstate analysis

The microstate analysis was performed using the Microstate EEGLAB Toolbox (Poulsen et al., 2018). Before running the microstate analysis, data from all sessions were concatenated to create one continuous epoch of around 20 minutes. Extracting microstates on the aggregated data improves the quality of the decomposition and facilitates a direct comparison of microstate dynamics across sessions.

Detailed descriptions of microstate extraction can be found in previous publications (Wackermann et al., 1993; Michel and Koenig, 2018; Mishra et al., 2020). Here we provide a brief overview. Microstate extraction is performed at global field power (GFP) peaks, defined as the standard deviation of the data across channels. We ran the analysis with a minimum peak distance of 10 ms and the number of GFP peaks was capped at 10% of the trial length. In our experience, the number of GFP peaks was always less than this upper limit. We used a modified k-means method for segmentation into 4 clusters. The number of clusters was chosen based on the Global Explained Variance (GEV) criterion (equation 2 below and Figure 1C_1_-C_3_). We noticed that the incremental change in the GEV saturated after 3-5 clusters. Moreover, cluster centers with similar spatial patterns started emerging after four clusters in most of the cases. A constant number of clusters across recording is useful for comparison purposes. Thus we chose to extract four clusters for all datasets. The segmentation process was repeated for 10 iterations and the iteration with the highest GEV was used for the segmentation. The randomness is caused by the random selection of cluster centers in the LFP signal to initiate the clustering process (Murray et al., 2008; Milz, 2016). The microstate maps extracted during this segmentation process were then backfitted on the data. The backfitting process means assigning a microstate at each time point based on the highest absolute correlation with all four microstate maps extracted in the previous step. This analysis was polarity invariant, which means that LFP data at two time points with similar spatial pattern but opposite polarity were labeled as the same microstate. Minimum duration of microstates was set to be 25 ms (Poulsen et al., 2018).

**Figure 1.**
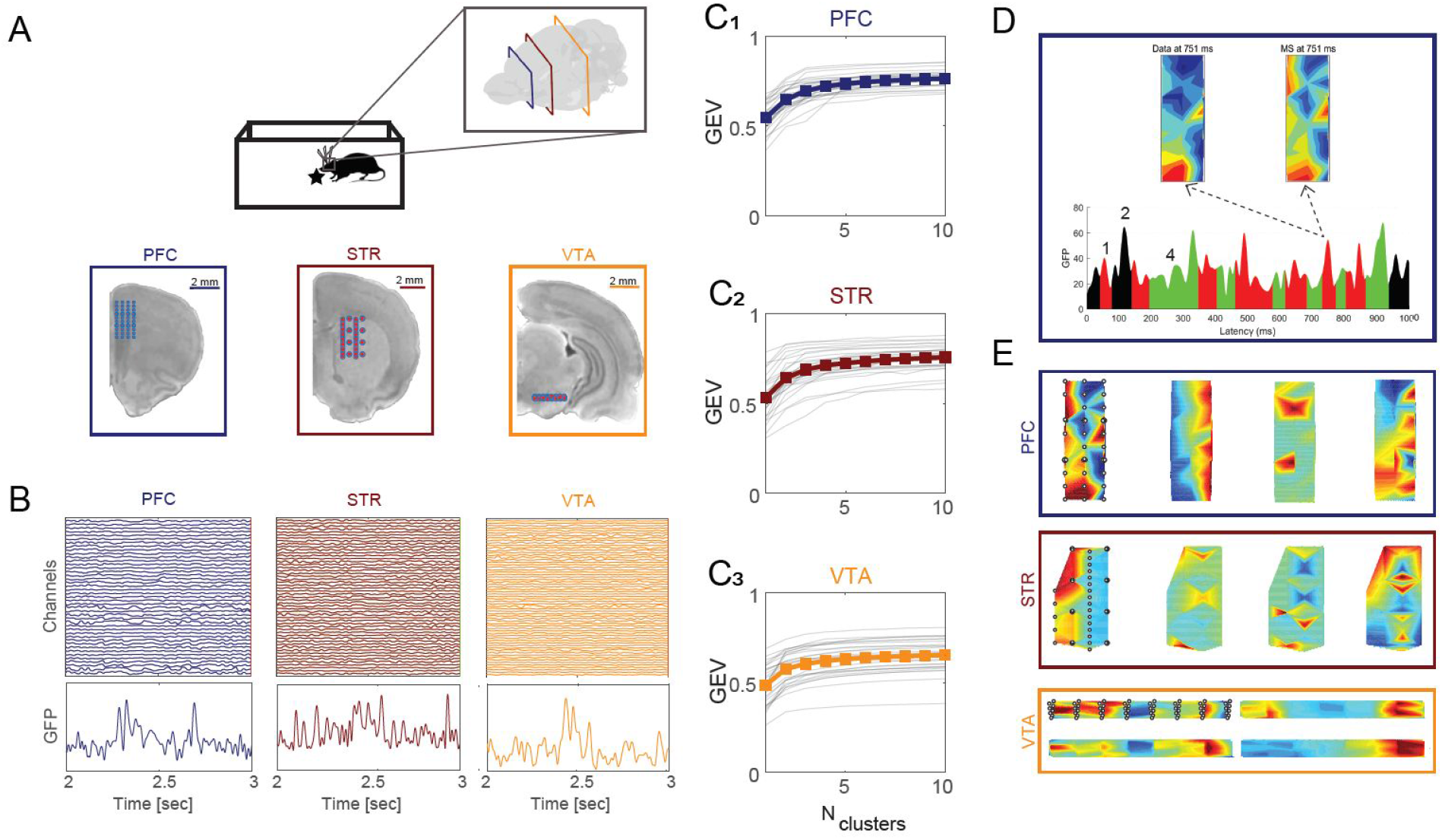
LFP signals recorded from multiple sensors can be represented by LFP microstates. **A,** Neural recordings were concurrently collected from three brain regions (PFC (navy), STR (maroon), VTA (orange) in awake behaving rats. Large-scale electrode arrays (red dots) covered a coronal plane in each brain area (slices show approximate antero-posterior locations, taken from the Sprague Dawley rat atlas of (Papp et al., 2014). **B,** LFP data excerpt from all channels recorded in each brain region (3×64, top) and global field power (GFP, bottom). Microstate (MS) maps were extracted based on the local peaks of the GFP, using a modified k-means clustering algorithm (Michel and Koenig, 2018; Poulsen et al., 2018; Mishra et al., 2020), see Methods for Details). **C_1_-C_3_,** Cumulative global explained variance saturated after 3-5 microstates in all three regions. Gray lines indicate results from individual datasets, thick lines indicate mean. **D,** The spatial patterns of LFP signals (top left) can be captured by a small number of MS maps (top right, e.g. MS1 at 750 ms. We focused on the four dominant microstates in each region (bottom, different colors). The current MS is determined by correlation between microstates and the current LFP. **E,** MS maps extracted from each region while the animal was freely moving in an open field environment (see A, top left). The left-most maps also show the electrode locations. Map values were spatially spline-interpolated between electrode locations.

### Microstate properties estimation

We quantified the temporal pattern of occurrence of microstates using the following five properties.

1. **Mean spatial correlation:** Defined as the mean Pearson correlation between LFP signals and their corresponding microstate map. We used the absolute value of the correlation for polarity-independent analyses.

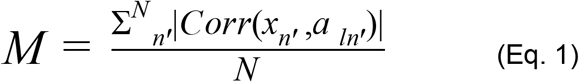 Where *x_n_* is the channel vector at time point *n*, *a_ln_* is the microstate map *l*, and *GFP_n_* is global field power, where *N* is the length of the signal.
2. **GEV:** Global explained variance is a measure of similarity between the data and the microstates. GEV for the n^th^ sample of the data is defined as,

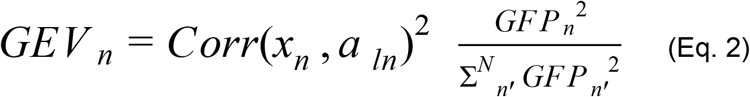
3. **Occurrence Rate:** The mean number of appearances of a microstate per second.
4. **Temporal Coverage:** The fraction of total time covered by all occurrences of a given microstate.
5. **State Duration:** Defined as the median duration of each microstate occurrence.

### Animal movement tracking and behavioral state determination

Animal movement tracking was performed using DeepLabCut (Mathis et al., 2018). We used this deep neural network-based method to track 5 body markers, i.e. snout, left ear, right ear, head center and tail start (Fig 4A_1_-A_2_). We also tracked the location of the object center, for sessions in which an object was present. This was done because the objects could be moved around slightly as the animals explored it. A moving median filter with a window length of 13 frames was used to post-process the movement tracking results. This improved accuracy by removing isolated video frames with incorrect location of body markers and objects.

In order to determine the video frames where the animal was not moving, a threshold on the speed of the animal was manually set based on visual inspection (see Fig.4D_2_).

We defined a radius around the object center, based on the size and shape of the object and determined separately for each object. For the extraction of frames where the rats were interacting with the object, the distance between the snout and the object center was computed. “Object interaction” was defined as frames in which this distance was less than the interaction radius. Two binary vectors, associated with two types of behavioral states - object interaction and movement - were upsampled to 1 kHz to match with the LFP data.

### Surrogate data for permutation test

In order to test the statistical significance of GEV by four microstates, and the effect on microstate properties by the change in behavioural states, we generated surrogate datasets in which the local temporal structure per channel was preserved, while the temporal alignment between channels was randomized. This was achieved in the LFP data by choosing a random point in time for each channel, and swapping the time-series around these time points. As opposed to a random shuffling of the signal in time and channel, this leaves the local temporal structure of the signal intact (Fig 3A-B). 100 surrogates were produced, and GEV were calculated for each surrogate to compare with the GEV of the empirical data. We also generated surrogate movement data using the same method. Again, 100 surrogates were produced. Microstate properties for each microstate were calculated for the surrogate movement data and compared against their empirical value to compute the significance (Fig 5C_1_-C_3_).

### Microstate activation cross-correlation and their power spectra

The ephys data were projected on the extracted microstates to generate microstate activation time-series. Cross-correlations were performed on all pairs (within and across regions) of these microstate activation time series, up to a maximum lag of ±2000 ms (using xcorr in MATLAB). The power spectra of the cross-correlograms were estimated by taking the square of the discrete Fourier transform (fft in MATLAB) divided by the length of cross-correlograms. To facilitate comparison and subsequent analyses, power spectra were z-normalized by taking the standard deviation over frequencies.

### Power-spectrum clustering

The cross-correlation power spectrum profiles were clustered using the k-means method. Squared Euclidean distance criteria was used as the distance metric for the clustering. The optimal number of clusters, determined using silhouette measure, was two. Clustering was performed separately for each dataset.

### Statistical analysis

Statistical analysis on microstate properties and their relationship with the behavioral states was performed without distributional assumptions, by using permutation tests. Statistical analysis on behavioral state durations and power spectra peak frequencies were performed using a two-sided Wilcoxon rank sum test.

## Results

We investigated the existence and dynamics of coordinated neural activity over three brain regions - prefrontal cortex (PFC), striatum (STR) and the ventral tegmental area (VTA) - in chronic recordings in behaving rats (N=4, Fig. 1A). Recordings were collected simultaneously using up to 192 electrodes (64 per region) in conjunction with video recordings. We used *microstate analysis*, an established techniques for detecting short-lived states of joint activation from the EEG literature (Lehmann, 1989; Michel and Koenig, 2018; Mishra et al., 2020) and deep-learning based video tracking (Mathis et al., 2018) to relate the occurrence of the detected *microstates* with ongoing behavioral states. We start by introducing the analysis technique to isolate microstates in multi-channel LFP data.

### Large-scale LFP signals are accounted for by a small set of microstates

Multiarray, multiarea LFP signals were acquired simultaneously (Fig. 1B) and subjected to microstate analysis separately per region. The microelectrode array geometry enabled the acquisition of neural signals from a wide area (0.15 −2 mm^2^). In each area, the spatial activity could be accounted for by a small number of microstates, i.e. joint activity patterns, whose residence time ranged between a few tens and a few hundreds of milliseconds (Fig. 1D, showing a sample from PFC). The segmentation of the LFP signal into microstates is based on spatial correlation of neural activity (for details see Methods). Qualitatively, this becomes apparent when the LFP data and corresponding microstate are visualized together (Fig. 1D, top, quantified in Table 1). The first four microstates accounted for 68% of the variance in the example shown (see Table 1 for region specific details). The exact number of significant microstates from a clustering analysis can be difficult to ascertain (Murray et al., 2008; Michel and Koenig, 2018); we chose to use four microstates per region for each recording day, based on the total amount of explained variance (Fig. 1C_1_-C_3_), to facilitate analyses across days. Although other choices may yield slightly different results, our experience is that the patterns of findings are robust to the exact number of microstates.

**Table 1:** Summary of microstate properties.

|  | **PFC** | **STR** | **VTA** |
| --- | --- | --- | --- |
| **Mean Spatial Corr** | 0.55 ± 0.08 | 0.54 ± 0.09 | 0.49 ± 0.08 |
| **Temporal Coverage** | 0.23 ± 0.14 | 0.23 ± 0.16 | 0.26 ± 0.10 |
| **Occurrence Rate (per sec)** | 3.65 ± 1.40 | 3.11 ± 1.38 | 4.08 ± 1.15 |
| **State Duration (ms)** | 68.04 ± 36.08 | 70.30 ± 26.57 | 65.41 ± 11.58 |
| **GEV (from top 4 MS)** | 0.71 ± 0.06 | 0.71 ± 0.09 | 0.64 ± 0.10 |

Notes. Properties are averaged across all recordings ( $N_{\text{recordings}}=27$ , $N_{\text{rats}}=4$ , mean ± S.D.).

Microstate topographies exhibited a mixture of distributed spatial features (Fig. 1E) (e.g. see MS_1_ (left) for STR) and restricted local activity (e.g. MS_2_ (middle-left) for STR). Importantly, electrodes in the array were individually inspected for spurious recordings, e.g. caused by unreasonably high variance or large differences in impedance (see Methods for details). Electrodes with excessive variance or artifacts were excluded from subsequent analysis, ensuring that the microstates reflected spatiotemporally coherent activation patterns based on well-sampled neural activity.

### Microstate maps are robust over multiple days

We repeated the experiment in the same animals over multiple days in order to determine the stability of microstate maps over time. The microstate analysis was done separately on each day’s recording (between 5 and 9 days per animal). We found that spatial correlations of microstate maps across days were high (Fig. 2), indicating that many of the maps recurred on different days. Other microstates were less stable and were identified on some but not all days (e.g. MS3/4 (blue/green) in PFC, Fig. 2A1). It is possible that these “missing” microstates were sorted into smaller clusters that were excluded from the primary analysis due to lower signal-to-noise or infrequent occurrences.

**Figure 2.**
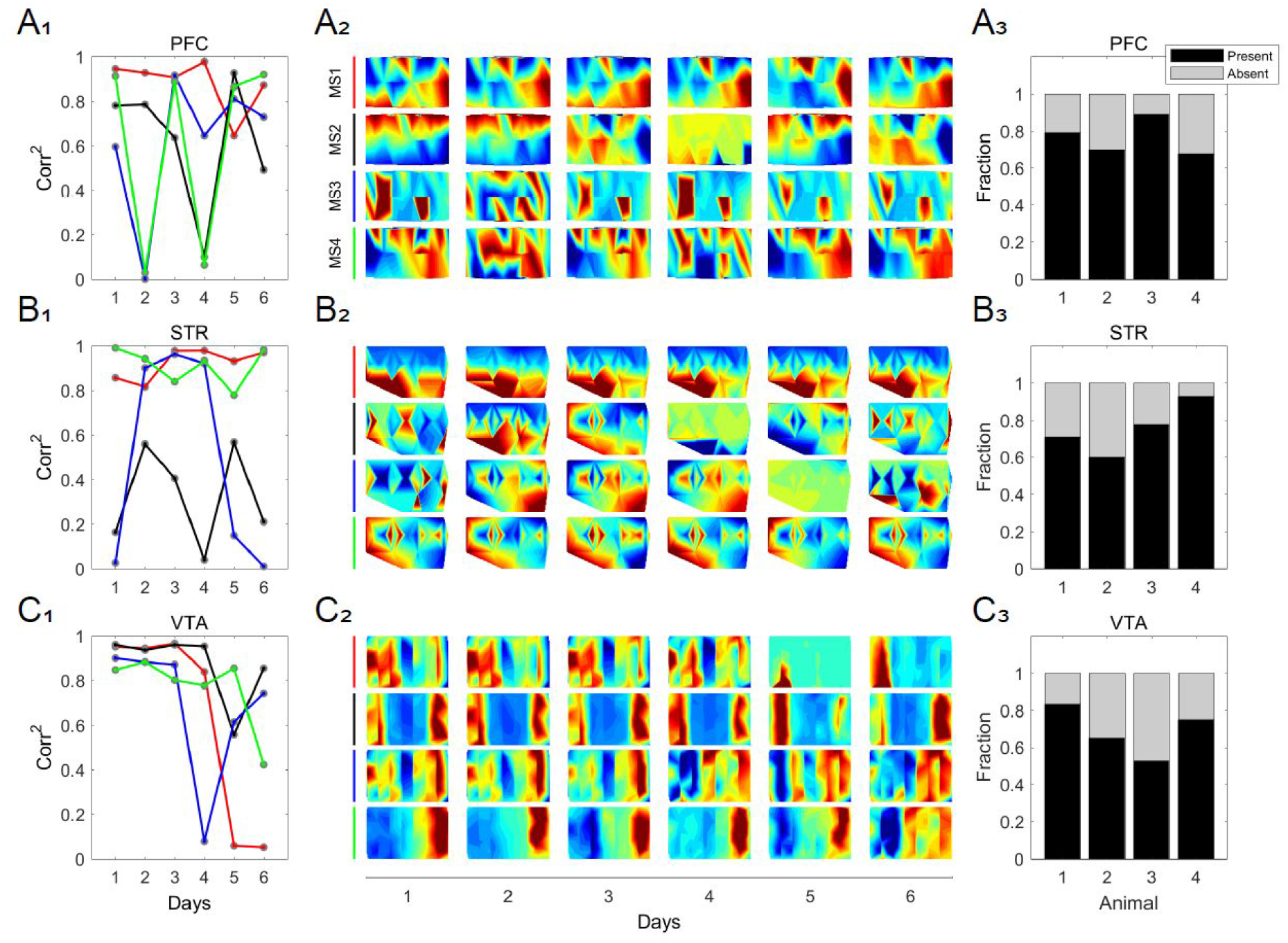
MS maps are reproducible across different days. Simultaneous LFP signals were recorded on 6 different days (each day with a new object) in PFC, STR and VTA. MS maps from each region on the first day were chosen as reference maps. Maps are rotated counter-clockwise by 90 degrees for PFC and STR and rescaled for visualization purposes. **A_1_,** In the PFC, the squared correlations between the four maps on each day and average microstate maps were computed, and are shown with red, black, blue and green lines for microstates 1 to 4, respectively. Lower correlation values on certain days can be loosely interpreted as the absence of the MS on that day (in reference to the average MS). **A_2_,** MS maps are reordered based on their best match (Panel A_1_). Most MS maps retained the spatial features on each recording day. (A_1_ and A_2_ are from animal 1 in panel A_3_) **A_3_,** The fraction of maps that were repeated over time for each of four animals. **B, C,** These rows show the same results as described above but for data from the striatum and VTA.

To assess the presence of microstate maps on experiment repeats, we averaged all highly similar maps (per region and per animal) together as their first principal component. We refer to these maps as subject average microstate maps (MS_sa_). Squared spatial correlations between the subject average microstate maps and microstate maps on each experiment day were computed as an indicator of MS recurrence (see Fig 2A_1_, 2B_1_, 2C_1_ for an example from one animal). We used a threshold of 0.5 (squared spatial correlation) to determine that a microstate repeated, although exploration of a range of threshold values (0.4 to 0.7) yielded qualitatively similar results. In all three regions, the same set of microstates, within the same animal, showed a stronger repeatability over separate recording days (Fig 2A_3_, 2B_3_ and 2C_3_).

### Spatiotemporal LFP structure underlies the microstates maps’ explanatory power

The previous analyses demonstrate that microstates are rather stable over multiple days. We next investigated how much variance in the total signal is explained by the microstates. We quantified this using the global explained variance (GEV), which reflects the correlation between the signal at each time point and the microstate maps. Using four microstates per region accounted for 71±6 %,71±9 % and 64±10 % (mean ± S.D.) of the global variance in PFC, STR, and VTA respectively. We tested whether these GEV values were statistically significant by generating a null hypothesis distribution, generated by shuffling the data in a way that preserved the local temporal structure but distorted global temporal structure (see Methods, Fig. 3A-B for illustration). Microstate analysis was rerun 100 times with different random surrogate datasets, and subsequently the GEV was recomputed.

**Figure 3.**
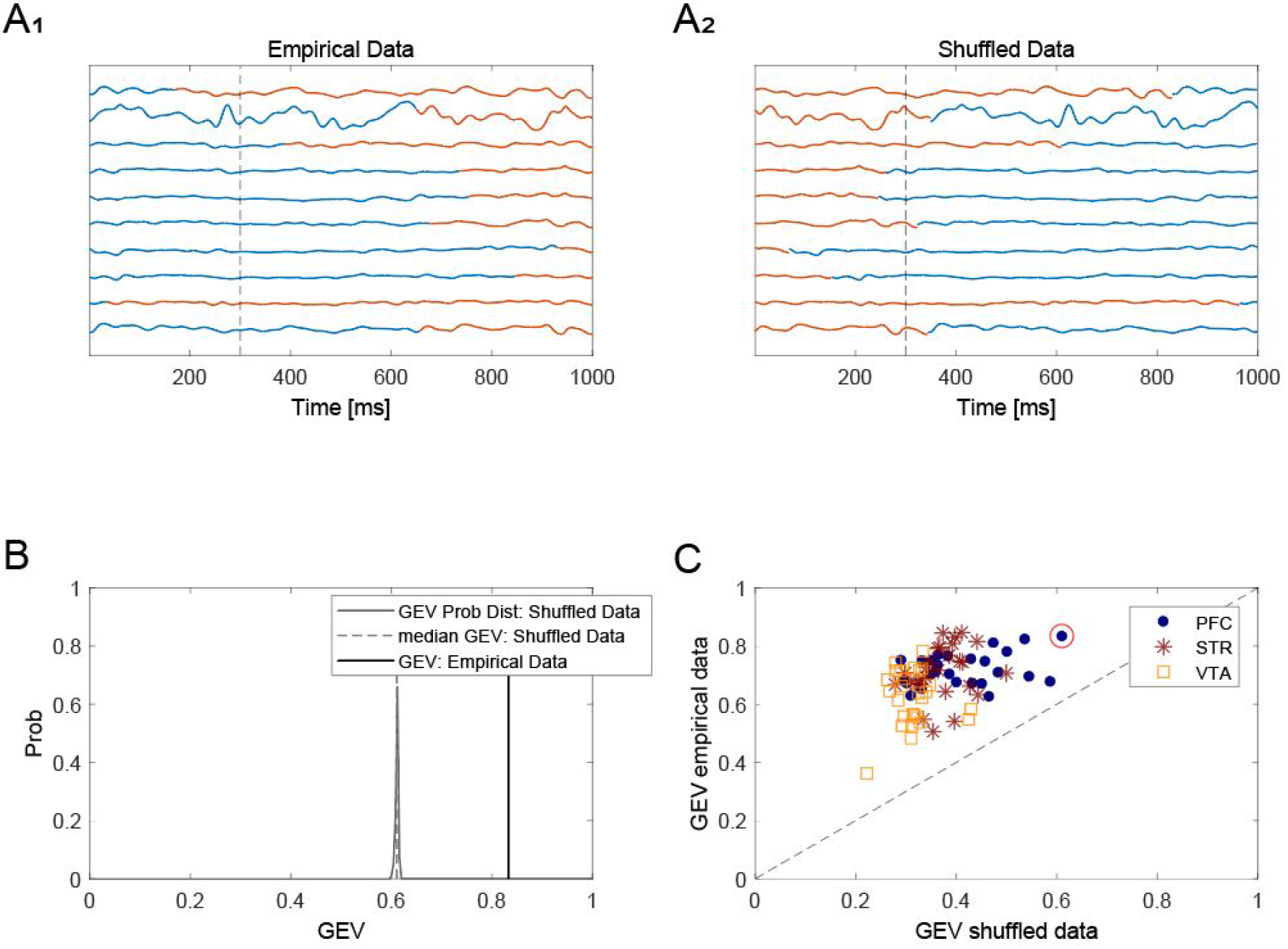
MS maps represent the spatiotemporal structure of LFP activity. **A_1_,** Surrogate LFP data sets with conserved local structure were created by pivoting the data around a random time point (independently selected per trial, blue = before pivot point, red = after pivot point). **A_2_,** The data were swapped around these pivot points to generate a surrogate dataset. **B,** MS analysis was performed on these surrogate datasets, and GEV was computed. The corresponding GEV distribution for one of the recordings (red circle in **C**) showed a significant decrease in GEV in the surrogate data, indicating that the original temporal alignment between channels is underlying the microstate maps explanatory value. **C,** GEVs of the empirical data were always higher than the corresponding median GEV from surrogate datasets in all three brain regions, showing statistical significance of the MS results.

We found that this removal of spatiotemporal LFP patterns in the surrogate data resulted in a significant drop in GEV (p<0.001) (Fig. 3C). The GEV of the shuffled data was only 41±9%, 37±5% and 31±4% in comparison to ~70% for the original data (see above), corresponding to a significant reduction by 43%, 47% and 51% (p<0.001, permutation test) in PFC, STR, and VTA respectively. GEV values differed considerably across regions (Fig. 3C), but in all cases using the spatial surrogates dropped the GEV significantly. In conclusion, microstates capture repeatedly occurring spatiotemporal activation patterns, a small number of which are sufficient to explain up to ~80% of the total LFP variance.

### Linking microstates to behavioral states

During the recordings, animals interacted with a novel or a familiar object in the arena. We subdivided the recording session along two behavioral dimensions: (1) “Movement-related” i.e. stationarity or in motion; and (2) “Object exploration-related” i.e. interacting or not interacting with the object. Tracking of the animal and the object was performed using DeepLabCut (v.2.1, Mathis et al., 2018) and each time point was assigned a binary value for each of the two dimensions based on empirically determined thresholds (Fig. 4, see Methods for details). Each recording day comprised four sessions (Fig. 4B); movement tracking was performed separately for each session and then later pooled.

**Figure 4.**
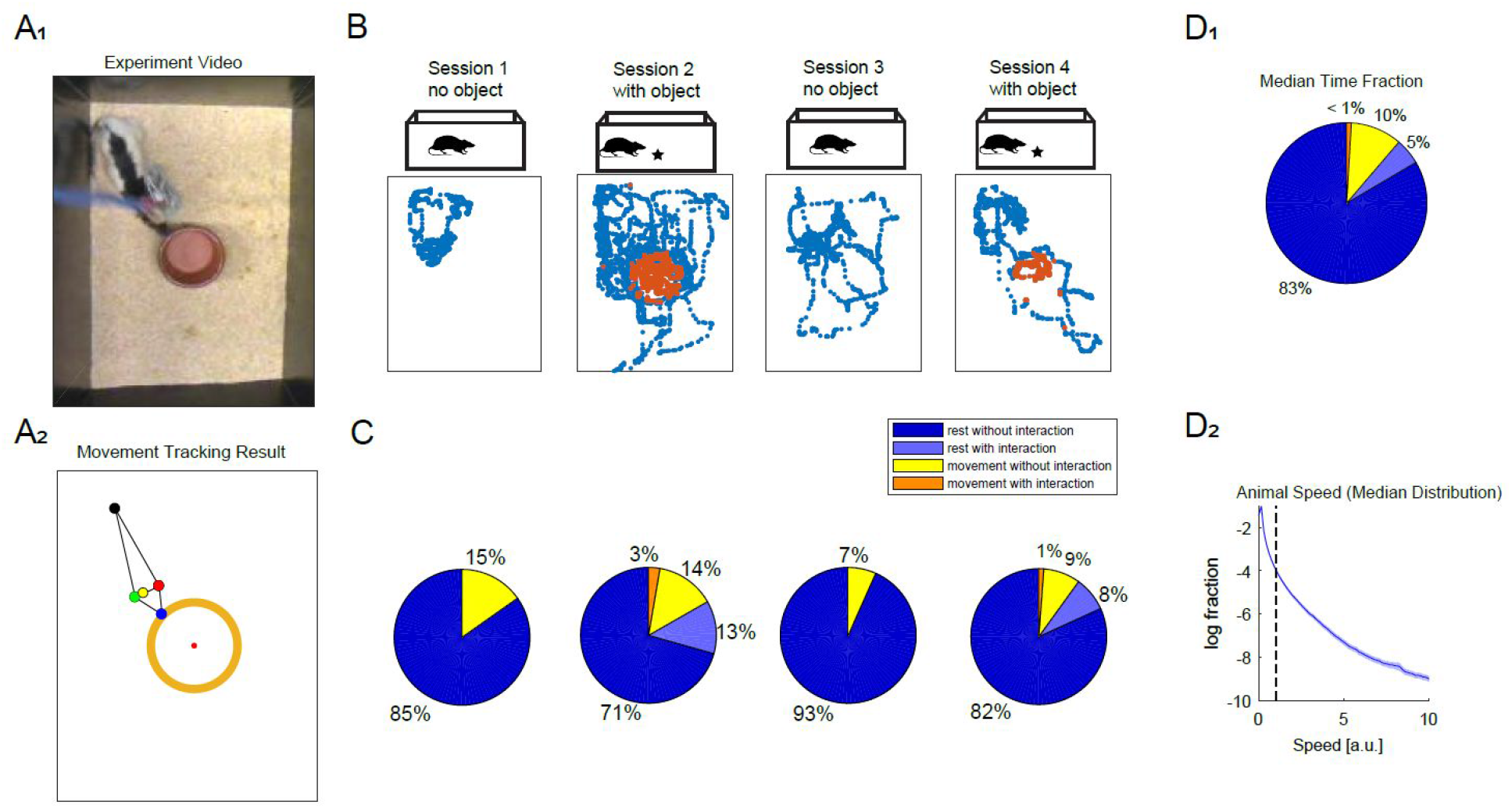
Automated movement tracking to segregate behavioural states. **A_1_,** An example video frame during the experiment. **A_2_,** Animal movement is accurately tracked using DeepLabCut, shown here using the same frame as in A_1_. Animal movement was tracked using five body markers (blue, black, red, green and yellow dots represent, respectively, snout, tail start, left ear, right ear and head center). **B,** Each experimental recording consisted of four sessions (~5 min each session). Sessions 1 and 3 were open field recordings, while session 2 included a novel object (different each day) in the middle of the box. In session 4, the same object was present. Rats spend less time interacting with familiar objects than novel objects (each dot represents snout location in each frame; red dots show time points labeled as interaction). A small number of erroneous object locations were filtered out prior to analysis. **C,** The distribution of temporal coverage of each behavioural state (rest, movement, object interaction) across sessions was dominated by periods of rest. Object interaction can take place during rest as well as during movement, represented as an overlap in behavioral states. **D_1_,** Median temporal coverage from all recordings captures the trends over all sessions, recordings and animals (N=4). **D_2_,** Median distribution of animal speed movement from all the recordings. The dashed line shows the motion speed threshold separating resting from movement.

Animals spent most of their time in the stationary state (88 ± 5%, median + S.D.) followed by movement state (12 ± 5%). Next, we classified these two behavioral states further into object interaction and no object interaction and found that the fraction of time spent on object interaction mostly took place during stationary state (5 ± 4%) compared to interaction during moving state (0.9 ± 0.7%) (Fig. 4C).

We next examined the evolution of behavioral states as time progressed. First, we compared the sessions in which the object was not present (sessions 1 and 3). We found that the animals spent more time in a stationary state in the later session (session 3, 93 ± 6%) in comparison to the earlier session (session 1, 85 ± 8%, p<<0.05, two-sided Wilcoxon signed ranks test). Next, we compared the time spent in stationary state when the object was introduced (sessions 2 and 4). We again found that animals spent more time in the stationary state in the later session (90 ± 5%) compared to the earlier session (83 ± 6%, p<<0.05, two-sided Wilcoxon signed ranks test). A novel object was first presented in session 2 and the same object was again presented in session 4. The fraction of time spent on object exploration in session 4 (9 ± 12%) was significantly less (p<0.05, two-sided Wilcoxon signed ranks test) than in session 2 (15 ± 12%). There were no trends along experiment repeats, indicating that the animals remained interested in the novel objects each day.

### Microstate dynamics were modulated by behavioural state

We quantified three microstate properties during each behavioral state — *Occurrence Rate, Temporal Coverage* and *State Duration* (see Methods for definitions) — and reported their summaries in Table 1. We separated these features for each microstate and brain region, and for the behavioral states ‘stationary’ and ‘movement’. All three microstate properties in all three brain regions were modulated by behavioral state (Fig. 5A shows an example from one recording in PFC). Furthermore, these modulations were consistent across the different recording days (e.g. Fig. 5B_1_-5B_3_ for one animal in PFC). Microstates followed the same direction of state-related changes in most of the repeats (PFC 76 ± 13%, STR 82 ± 14%, VTA 82 ± 15%), indicating a systematic relationship between microstates properties and behavioral states.

**Figure 5.**
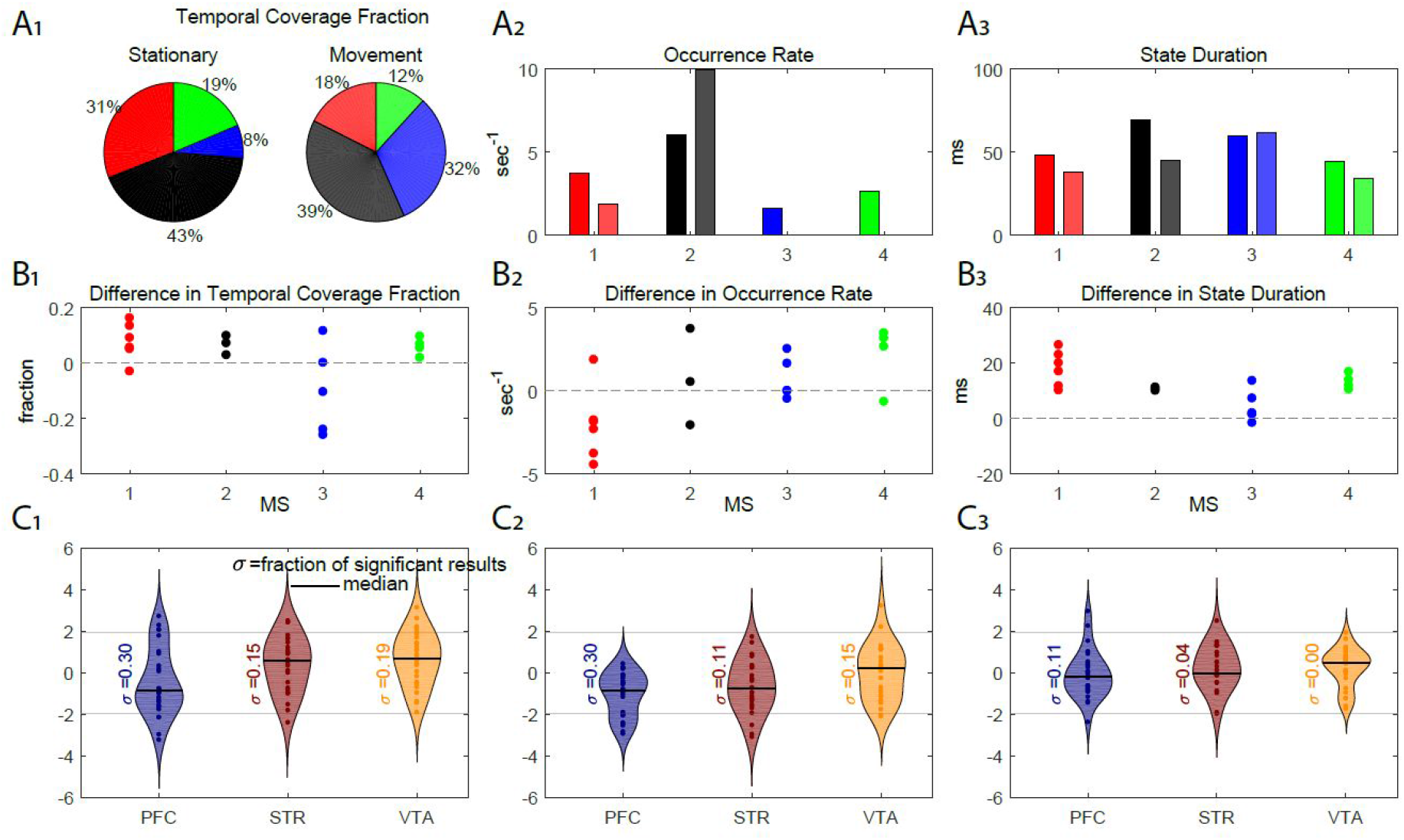
MS parameters are affected by behavioural state. **A,** example results from the recording in PFC of one rat on one recording day. **A_1_,** Temporal coverage fraction for all four MS differed in the two behavioral states (statistics in 5C_1_) **A_2_,** Occurrence rate during the two behavioral states (solid bars on the left refer to stationary periods, and lighter bars on the right refer to movement periods; the four numbers refer to the different MS). **A_3_,** Similarly, MS durations were modulated by behavioral state. **B_1_-B_3_,** The modulation of MS properties by behavioural state (expressed as stationary minus movement) was consistently in the same direction (PFC 76 ± 13%) across 6 different recording days (each dot corresponds to a day; only microstates with r^2^ > 0.5 repeatability are shown here). **C_1_-C_3_,** Similar to panel B but for data pooled across 27 novel objects in four rats. Significance was assessed by comparison against surrogate datasets (thin horizontal lines indicate a significance threshold of z = ±1.96). There was a high fraction of significant results for all three MS properties in all three brain regions.

We assessed the statistical significance of the behavioral state-dependency by generating a null hypothesis distribution based on surrogate behavioral states, obtained by random temporal cuts of the time series of behavioral labels (Fig. 3A and Methods). *Temporal Coverage* changed significantly between behavioral states, as all three regions showed a high fraction of significant changes (|z|>1.96) exceeding 0.05 (fraction of significant tests: PFC: 0.76; STR: 0.79; VTA: 0.66). *Occurrence Rate* showed a similar trend. The changes in the *Occurrence Rate* were also significant in all three regions (PFC: 0.35; STR: 0.44; VTA: 0.52). *State Duration* was less affected by behavior (PFC: 0.22; STR: 0.23; VTA: 0.08) (Fig. 5C). We also repeated these analyses for object exploration-related behavioral states, and observed qualitatively similar results.

In summary, temporal dynamics of microstates were significantly affected by changes in behavioral state. The most significant changes were observed for *Temporal Coverage.*

### Relationship of microstate dynamics across brain regions

We speculated that if microstates reflect a mesoscale organizational principle, then functional connectivity across brain regions might manifest as correlated microstate temporal dynamics. To investigate this possibility, we computed cross-correlations across all pairs of microstate activation time series, defined as the projection of sensor level signals at each time point onto the microstate basis vectors, in the three regions (144 cross-correlation pairs in total, out of which 12 are autocorrelations and 66 are inter-regional; omitting pairs that only differ by sequence) (Fig. 6A). All cross-correlation and auto-correlation curves followed a damped oscillatory nature with an average peak of 9 ±6 ms. The peak amplitude of cross-correlations varied substantially across microstate combinations and across animals (absolute correlation value 0.17 ± 0.07, mean ± S.D.).

**Figure 6.**
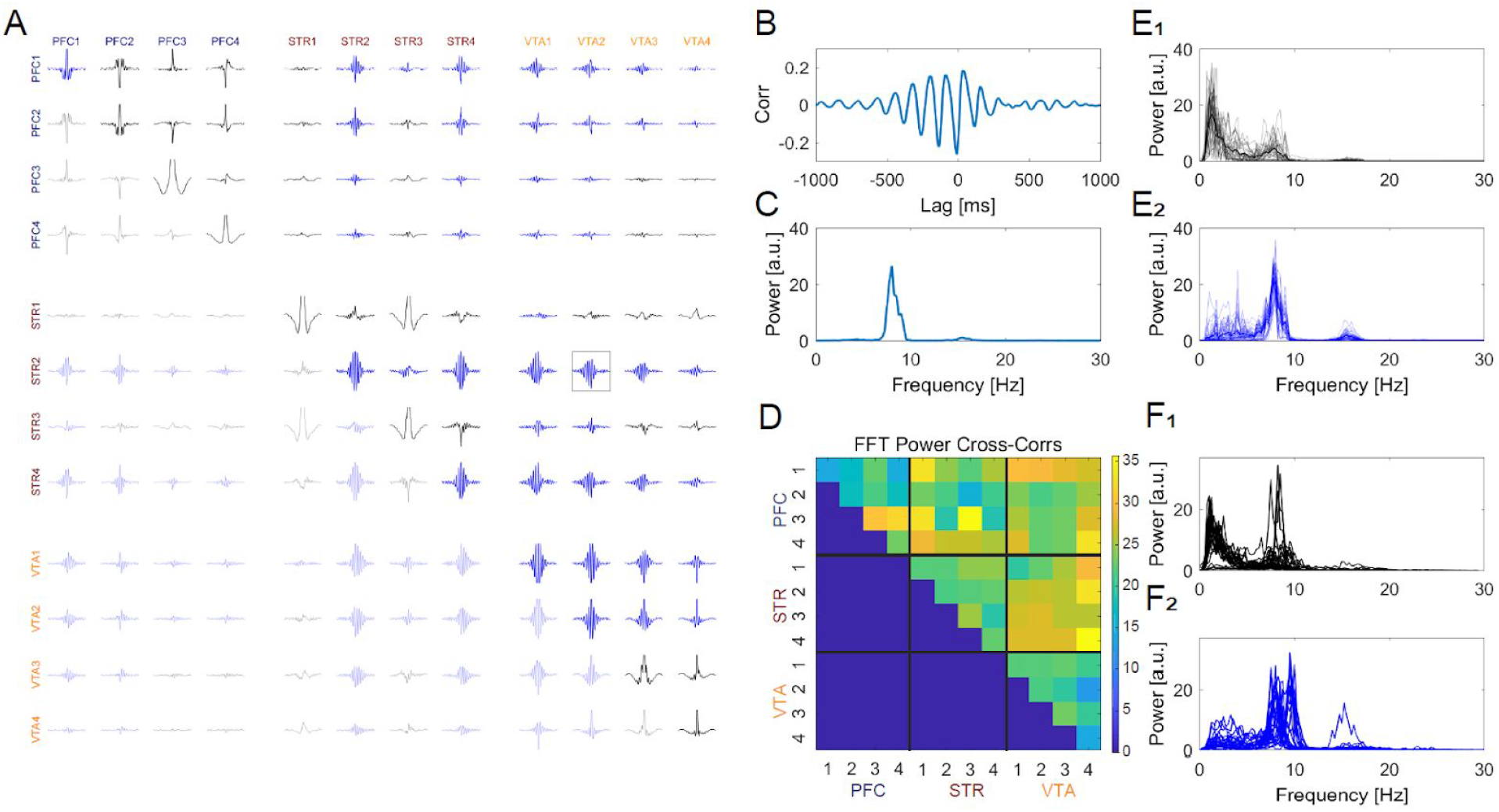
MS activation time-series are rhythmically correlated within and between brain areas. **A-E,** Illustrative results from one recording session. **A,** MS time-series cross-correlations show damped oscillatory dynamics both across MS within regions, and across regions. **B,** Example cross-correlation between activation time series of MS2 from STR and MS2 from VTA (this is a zoomed-in version of the boxed plot in Panel A). **C,** Z-normalized power spectrum of the cross-correlation shows a peak around 8 Hz. **D,** Z-normalized peak power from all combinations of microstate activation cross-correlations illustrates varying degrees of correlation within and between areas. **E_1_-E_2_** Z-normalized power spectra from all combinations of microstates within and between areas were clustered into two groups, one with a lower-frequency peak (~2-4 Hz, black lines, E_1_) and one with a higher-frequency peak (~8 Hz, blue lines, E_2_). Thick lines indicate cluster centers and thin lines are individual power spectra. **F_1_-F_2_,** Results from all recordings in all animals shows a consistent grouping of power spectra based on the frequency of the peak in power.

Remarkably, the cross-correlations exhibited robust oscillations, suggesting rhythmic patterns of temporal microstatic activation coordination across microstates and regions. The z-normalized power spectral profiles (Fig. 6C) contained a distinguishable set of peaks at different frequencies. Typically, one or multiple peaks were below 10 Hz, along with smaller peaks in higher frequencies (10-30 Hz). These higher-frequency peaks appeared to be harmonics of the lower frequency peaks, and we therefore focused on the lower frequencies. As expected from the cross-correlation profiles, peak power varied for different microstate combinations, suggesting the varying strength of cross-region interaction in different microstate combinations (Fig. 6D).

The power spectral profiles of the cross-correlations grouped into two clusters, based on k-means and the silhouette method (Rousseeuw, 1987). The first cluster had a dominant peak at 2-4 Hz while the second was dominated by peaks around 8-10 Hz (Fig. 6E_1_-6E_2_). These spectral profiles were consistent both within-animal (across experiment repeats) and across animals. All individual spectral profiles showed a consistent shape (Fig. 6F_1_-6F_2_).

In summary, microstate activations were temporally synchronized across brain regions both in slow (2-4 Hz) and theta/alpha (8-10 Hz) ranges. This suggests that mesoscopic-scale networks interact in a rhythmic fashion across the brain areas.

### Inter-regional interactions and behavioral states

Lastly, we investigated the possible behavioral relevance of the microstate activation correlations by separating them according to two behavioral states: object interaction and “long stationary periods”. We defined “long stationary period” as the trial segments where the animals remained stationary for longer than the 90th percentile of all stationary segment lengths (calculated separately for each trial, 2.3 ± 0.9 sec). While it is easier to distinguish moments of object interaction and no interaction, shorter segments of stationarity can be indistinguishable with movement preparation. Hence, in order to compare against the dynamics during the “true” stationarity, we chose to focus on longer stationary periods. Next, we defined dominant microstates during these two behavioral states in terms of their *Temporal Coverage* (Fig. 7A_1_-7A_3_). We focused on Temporal Coverage because it showed the most robust dependence on object exploration in all three regions. The cross-correlation power spectra during long resting periods dominated the 8-10 Hz peak whereas the power spectra during object interaction dominated the 2-4 Hz (Fig. 7B-C).

**Figure 7.**
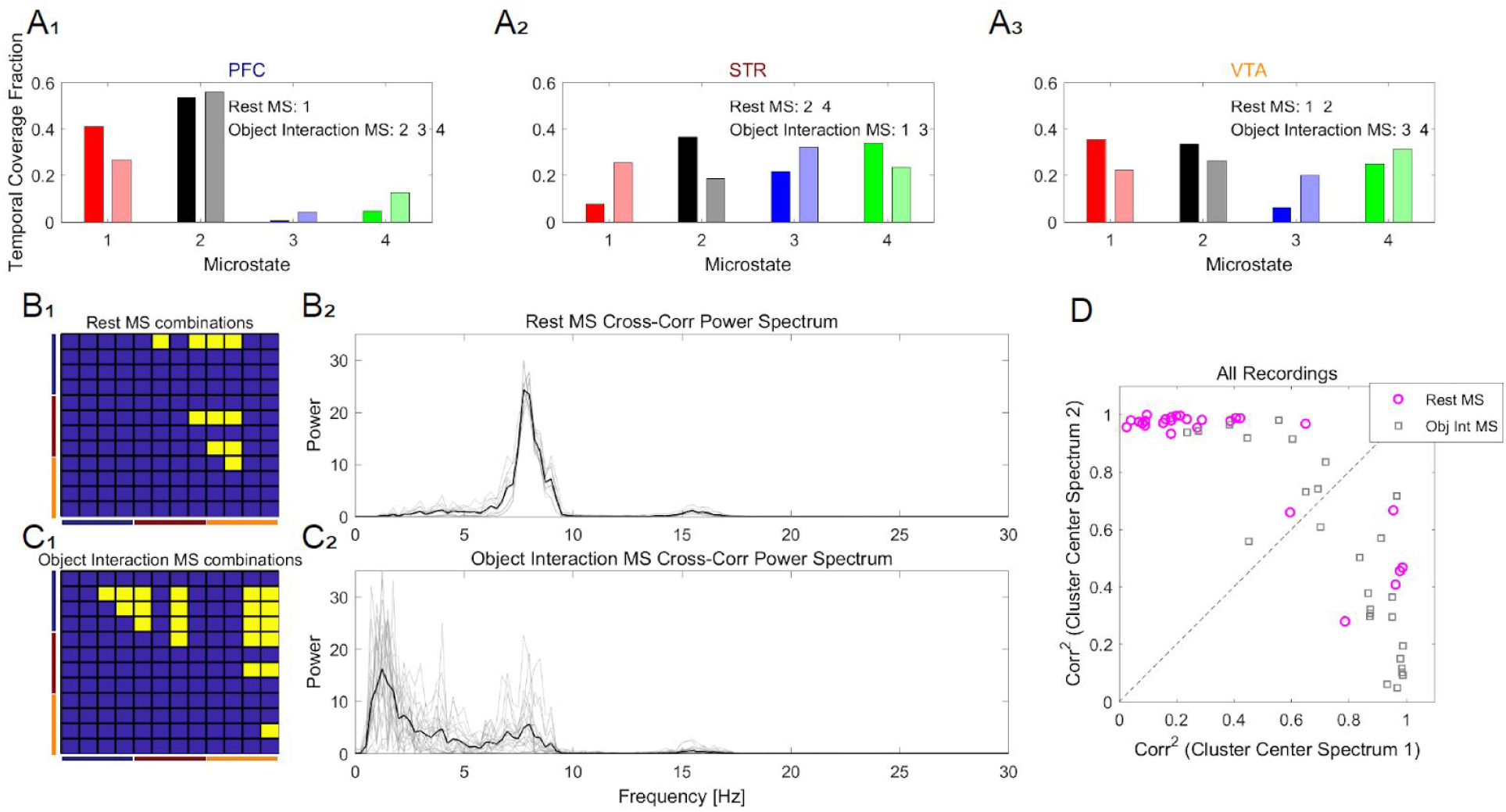
Power spectra of MS activation cross-correlations are grouped into two clusters that are differentially modulated by behavior. **A-C,** Illustrative data from one recording. **A_1_-A_3_,** Temporal coverage by each MS during longer resting periods (left bars) and object interaction (right bars shown in light colors) indicates the dominant MS during each behavioral state. **B_1_,** Dominant microstates (yellow) during longer stationary periods are shown across areas. All dominant MS were grouped for further cross-correlation spectrum analysis. **B_2_,** Power spectra of cross-correlation of relevant MS (shown in B_1_) during longer rest periods show a peak around 8 Hz (thick line shows the mean power spectrum). **C_1_,** Same as B_1_ for object interaction MS. **C_2_,** Cross-correlation power spectra of dominant MS (shown in yellow in C_1_) during object interaction show an earlier peak around 2-4 Hz (thick line shows the mean power spectrum). **D,** Mean power spectra from all recordings show higher affinity to one of the spectral cluster centers (shown in 6F_1_-F_2_), except very few spectra from object interaction MS combinations that show relatively greater similarity to the cluster more populated by rest MS.

**Figure 8.**
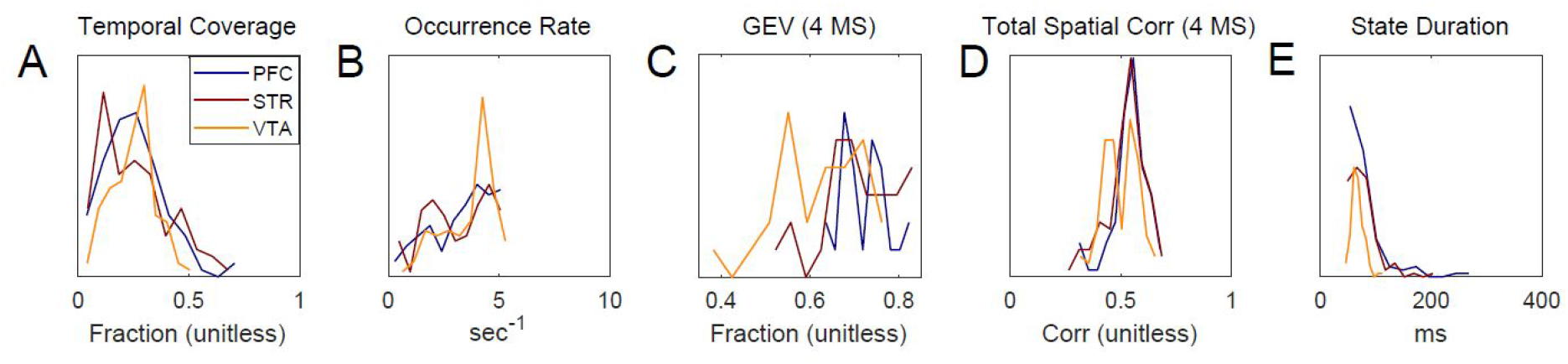
Distributions of MS properties in all three regions in all recording sessions provide insights on temporal and spatial features of MS (see also Table 1). **A,** Temporal coverage fraction shows temporal area covered by each microstate. **B,** Occurrence rate shows that MS in all three areas appeared roughly 3-4 times per second on average. **C**, The top four microstates exhibited a high Global explained variance (60-80%) in all three brain regions. **D,** The top four microstates exhibited high spatial correlation (Pearson) with the electrode topography (~0.5). **E,** State duration of MS followed power law distributions in PFC and STR, and had a characteristic peak of around 100 ms in the VTA.

Furthermore, power spectra associated with long resting periods were found to have a higher squared correlation with the 2-4 Hz cluster (r^2^ = 0.87 ± 0.22, mean + S.D.) compared to the 8-10 Hz cluster (r^2^ = 0.36 ± 0.32, p<<0.05, two-sided Wilcoxon signed ranks test) (Fig. 7D). On the other hand, the power spectra associated with object interaction correlated better with the higher peak frequency cluster centers (r^2^ = 0.76 ± 0.24) compared to the low peak frequency cluster (r^2^ = 0.50 ± 0.32, p<0.05, two-sided Wilcoxon signed ranks test) in terms of squared correlation (Fig. 7D). Thus, each behavioral state exhibited a distinct power spectrum profile. The two clusters of power spectral profiles (Fig. 6F) were modulated by object exploration and movement.

In summary, the microstate properties’ dependence on two contrasting behavioral states (object exploration and longer stationary periods) is supported by both the significant changes in microstate properties (Fig. 5) and the distinct nature of interaction patterns (cross correlation spectra) across regions. In fact, these two types of interaction indicators relate to the two behavioral states. This suggests the specificity of the underlying neural mechanism of the two behavioural states.

## Discussion

In this study, we extended a spatiotemporal analysis method developed for EEG data to characterize the dynamics of meso-scale signals (LFP) in multiple brain regions, and to search for states of activation and their interactions, during novelty-seeking in awake behaving rats. We identified microstates analogous to EEG microstates, which were physiologically interpretable and modulated by the behavioral state. This establishes a novel link between network-level activations and dynamics and concurrent behavioral states, and may facilitate bridging human EEG and rodent invasive research.

### “States” and scales in brain activity

The concept of a “brain state” appears in multiple domains of the neuroscience literature and spans multiple spatial scales and levels of analysis. For example, the “resting state” in fMRI (Raichle et al., 2001; Biswal et al., 1995) refers to coordination across widespread brain regions during rest. Large spatial scale states have also been defined based on temporally precise measurements such as EEG and MEG. Examples include defining entropy as the basis of states (Kannathal et al., 2005), using the embedded dimension in a state-space to define cortical synchrony (Carmeli et al., 2005), hidden markov model based definition of states (Baker et al., 2014; Hunyadi et al., 2019), and clustering-based approach of EEG data segmentation into quasi stable states (Lehmann et al., 1987b; Lehmann, 1989; Michel et al., 1999; Michel and Koenig, 2018).

At a finer spatial scale, neural states have been defined based on population spiking activity. For example, “up states” and “down states” in the context of depolarization and hyperpolarization (Steriade et al., 1993), the concept of “vocabulary” in population spike patterns (Luczak et al., 2009), propagating wave patterns based spatiotemporal structure in neural signals (Townsend and Gong, 2018), and in several other context of brain dynamics (Greenberg et al., 2008; Poulet and Petersen, 2008; Curto et al., 2009; Okun et al., 2010). Together, these studies show spatiotemporal coordination across neural populations activity. Travelling waves may be an example of mesoscopic states (Muller et al., 2018), as they have been observed across multiple spatial scales (Bahramisharif et al., 2013; Zanos et al., 2015; Alamia and VanRullen, 2019) and have been linked to a variety of cognitive functions.

Our approach was to adapt methods developed in the EEG literature to multichannel LFP recordings. Although it was in the present study not yet possible to directly compare our LFP microstates to human EEG microstates, many features of the identified LFP microstates appear to be consistent with a multitude of EEG microstate findings (Table 1) (Michel and Koenig, 2018), including the number of microstates to capture a significant amount of variance, their temporal characteristics, and relation to ongoing behavior. Interestingly, state durations in the range of 50-100 ms were observed in our data, in the EEG microstates literature (Michel et al., 1999), and in population spike recordings (Luczak et al., 2009). This hints at a possible common link underlying observable brain states across different spatial scales.

### Microstates and dimensionality reduction approaches

LFP microstates are identified by a data-driven dimensionality reduction approach that is based on long, continuous recordings and yet can be linked to transient behaviors. We found that four microstates can explain 60-80% of the LFP signals. This is overall consistent with other dimensionality reduction approaches, for example independent component analysis of LFP in the hippocampal CA1 (Makarov et al., 2010), principal component analysis in rat barrel cortex (Einevoll et al., 2007) and monkey PFC (Machens et al., 2010), demixed principal component analysis on the population activity in monkey PFC and rat orbitofrontal cortex (Kobak et al., 2016). It is remarkable that the conclusion of 3-6 large components persists over many different approaches. It seems that multi-channel LFP data occupy low-dimensional subspaces, and that different dimensionality-reduction methods identify different basis vectors to span a limited set of most relevant subspaces.

The maximization criteria of the different methods are important, however. For example, maximizing variance or non-Gaussianity do not necessarily lead to neurophysiologically interpretable components (de Cheveigné and Parra, 2014). On the other hand, the EEG microstate method uses modified k-means or hierarchical clustering of the data to produce basis vectors, and are thus easier to interpret physiologically and cognitively (Khanna et al., 2015; Michel and Koenig, 2018).

### LFP microstates exhibit both stability and variability over time

The reproducibility of the microstate maps across 5-9 days suggests that the microstates are stable over time within animals, perhaps reflecting the activation of circuits that are anatomically patterned. On the other hand, some microstate maps were not in the top four components in all recording sessions. It is possible that these maps had lower signal-to-noise and thus were relegated to lower-ranked microstates, or that these maps reflected state-specific functional networks that were less constrained by stable anatomical connectivity. On the other hand, the reproducible microstates were modulated by the behavioral states of the animals, indicating that their patterns of occurrence are not simply hard-wired. Overall, we find that LFP microstates are a promising tool to understand the activity dynamics underlying cognition.

### LFP microstates reflect mesoscale rhythmic coordination across brain regions

We identified a novel manifestation of rhythmic synchronization between mesoscopic-scale networks in spatially distinct brain regions. Remarkably, the interactions exhibited two prominent peaks in the lower frequency spectrum that mapped onto different aspects of behavior. The ~2 Hz peak was more strongly modulated by interacting with the novel object whereas the ~8 Hz peak was more strongly modulated by movement. This suggests that partly overlapping networks utilize spectral multiplexing to coordinate interregional interaction during novelty using two distinct routes of mesoscopic information exchange (Akam and Kullmann, 2014).

The PFC is crucial for higher cognition (Uylings et al., 2003; Laubach et al., 2018), and is involved in learning and attention (Passetti et al., 2002), decision making (De Bruin et al., 2000; Sul et al., 2010), working memory (Bekinschtein and Weisstaub, 2014), inhibitory response control (Brown and Bowman, 2002; Hardung et al., 2017), and other executive functions. The PFC also plays a crucial role in communication with other brain regions to execute many brain functions. This includes category learning through enhanced functional connectivity (Antzoulatos and Miller, 2014), encoding the reward information in the beta band (Zhang et al., 2016), among others (Feingold et al., 2015; Pasupathy and Miller, 2005; Gao et al., 2017).

Interactions amongst the PFC, VTA, and striatum are also crucial, particularly for learning and working memory, through dopaminergic pathways (Bentivoglio and Morelli, 2005; Björklund and Dunnett, 2007; Zahrt et al., 1997; Peters et al., 2004; Fujisawa and Buzsáki, 2011; Ott et al., 2014; Karreman and Moghaddam, 1996; Gao et al., 2007). It remains unclear whether the rhythmic coordination reflects long-range excitation or inhibition, however, the consistent structure of these microstate activation correlation profiles suggests that they play a meaningful role in relation to behavior.

### Implications for understanding novelty seeking

The experimental paradigm used here is a variant of the commonly used object recognition test (Aggleton, 1985). Novelty seeking has been shown to evoke exploratory behavior (Berlyne, 1950). The reduction of exploratory behavior during the repetition of a previously-novel object is taken as a measure of single-trial memory formation. The role and interplay of PFC, STR and VTA is well known for memory and motivation (Karreman and Moghaddam, 1996; Palmiter, 2008; Liljeholm and O’Doherty, 2012; Antzoulatos and Miller, 2014; Bekinschtein and Weisstaub, 2014; Morales and Margolis, 2017). The access to these brain regions makes this experimental paradigm suitable for exploring the neural states underlying the novelty seeking behavior. Moreover, the prolonged stationary behavioral state, as the trials progressed, allowed us to investigate and compare the neural activity during novelty seeking against the “background activity”. In this study, we propose a role for mesoscale microstates in novelty seeking behavior, which can be attributed to the activation of different networks during novelty processing.

### Limitations and Future Directions

While this study lays the foundation of LFP microstates, it also presents a few limitations, in particular their interpretation, spatial reach, and origin of interregional coordination.

First, the interpretation of the LFP microstates is not self-evident: one exciting possibility would be that LFP microstates reflect building blocks of brain-wide neural activity (in the EEG literature even termed “atoms of thought”; (Lehmann, 1990) or instead as a mesoscopic equivalent of default mode networks (Biswal et al., 1995). Second, the “true” number of microstates and their overall shape (the spatial structure) during different sensory stimulation and spontaneous conditions is difficult to determine even with multi-electrode arrays. Future studies should use larger arrays to address the spatial reach of microstates. Third, while we were able to connect the microstate approach with the previously known results of inter-regional communication in an interpretational sense, by establishing the cross region interaction using the microstate activation and link them with behavioral states, their relation to other manifestations of connectivity remain to be explored.

We aim to answer these questions in future studies using larger, multiscale electrodes. While we extended microstate analysis here to LFPs, the next important challenge lies in combining EEG and LFP measurements.

## Acknowledgements

The authors would like to thank Dr. Paul Anderson for his contribution in the surgeries and data collection process. MXC is funded by an ERC-Stg (638589) awarded to MXC. AM is funded by a joint doctoral grant awarded to MXC and BE by the Donders Center of Neuroscience. BE was supported by a European Commission’s Marie Curie Grant (660328), a NWO VIDI Grant (016.189.052) and a NWO ALW Open Grant (ALWOP.346).

## References Bibliography

Aggleton JP (1985) One-Trial Object Recognition by Rats. The Quarterly Journal of Experimental Psychology Section B 37:279–294.

Akam T, Kullmann DM (2014) Oscillatory multiplexing of population codes for selective communication in the mammalian brain. Nat Rev Neurosci 15:111–122.

Alamia A, VanRullen R (2019) Alpha oscillations and traveling waves: Signatures of predictive coding? PLoS Biol 17:e3000487.

Alishbayli A, Tichelaar JG, Gorska U, Cohen MX, Englitz B (2019) The asynchronous state’s relation to large-scale potentials in cortex. J Neurophysiol 122:2206–2219.

Antzoulatos EG, Miller EK (2014) Increases in functional connectivity between prefrontal cortex and striatum during category learning. Neuron 83:216–225.

Arieli A, Shoham D, Hildesheim R, Grinvald A (1995) Coherent spatiotemporal patterns of ongoing activity revealed by real-time optical imaging coupled with single-unit recording in the cat visual cortex. J Neurophysiol 73:2072–2093.

Bahramisharif A, van Gerven MAJ, Aarnoutse EJ, Mercier MR, Schwartz TH, Foxe JJ, Ramsey NF, Jensen O (2013) Propagating neocortical gamma bursts are coordinated by traveling alpha waves. J Neurosci 33:18849–18854.

Baker AP, Brookes MJ, Rezek IA, Smith SM, Behrens T, Probert Smith PJ, Woolrich M (2014) Fast transient networks in spontaneous human brain activity. elife 3:e01867.

Battaglia FP, Sutherland GR, McNaughton BL (2004) Hippocampal sharp wave bursts coincide with neocortical “up-state” transitions. Learn Mem 11:697–704.

Bekinschtein P, Weisstaub N (2014) Role of PFC during retrieval of recognition memory in rodents. J Physiol Paris 108:252–255.

Bentivoglio M, Morelli M (2005) Chapter I The organization and circuits of mesencephalic dopaminergic neurons and the distribution of dopamine receptors in the brain. In: Dopamine, pp 1–107 Handbook of chemical neuroanatomy. Elsevier.

Berlyne DE (1950) Novelty and curiosity as determinants of exploratory behaviour1. British Journal of Psychology General Section 41:68–80.

Biswal B, Yetkin FZ, Haughton VM, Hyde JS (1995) Functional connectivity in the motor cortex of resting human brain using echo-planar MRI. Magn Reson Med 34:537–541.

Björklund A, Dunnett SB (2007) Dopamine neuron systems in the brain: an update. Trends Neurosci 30:194–202.

Britz J, Díaz Hernàndez L, Ro T, Michel CM (2014) EEG-microstate dependent emergence of perceptual awareness. Front Behav Neurosci 8:163.

Britz J, Michel CM (2011) State-dependent visual processing. Front Psychol 2:370.

Brown VJ, Bowman EM (2002) Rodent models of prefrontal cortical function. Trends Neurosci 25:340–343.

Carmeli C, Knyazeva MG, Innocenti GM, De Feo O (2005) Assessment of EEG synchronization based on state-space analysis. Neuroimage 25:339–354.

Chiu C, Weliky M (2001) Spontaneous activity in developing ferret visual cortex in vivo. J Neurosci 21:8906–8914.

Curto C, Sakata S, Marguet S, Itskov V, Harris KD (2009) A simple model of cortical dynamics explains variability and state dependence of sensory responses in urethane-anesthetized auditory cortex. J Neurosci 29:10600–10612.

de Cheveigné A, Parra LC (2014) Joint decorrelation, a versatile tool for multichannel data analysis. Neuroimage 98:487–505.

Delorme A, Makeig S (2004) EEGLAB: an open source toolbox for analysis of single-trial EEG dynamics including independent component analysis. J Neurosci Methods 134:9–21.

De Bruin JPC, Feenstra MGP, Broersen LM, Van Leeuwen M, Arens C, De Vries S, Joosten RNJMA (2000) Role of the prefrontal cortex of the rat in learning and decision making: effects of transient inactivation. In: Cognition, emotion and autonomic responses: The integrative role of the prefrontal cortex and limbic structures, pp 103–113 Progress in brain research. Elsevier.

Dierks T, Jelic V, Julin P, Maurer K, Wahlund LO, Almkvist O, Strik WK, Winblad B (1997) EEG-microstates in mild memory impairment and Alzheimer’s disease: possible association with disturbed information processing. J Neural Transm 104:483–495.

Einevoll GT, Pettersen KH, Devor A, Ulbert I, Halgren E, Dale AM (2007) Laminar population analysis: estimating firing rates and evoked synaptic activity from multielectrode recordings in rat barrel cortex. J Neurophysiol 97:2174–2190.

Feingold J, Gibson DJ, DePasquale B, Graybiel AM (2015) Bursts of beta oscillation differentiate postperformance activity in the striatum and motor cortex of monkeys performing movement tasks. Proc Natl Acad Sci USA 112:13687–13692.

Fiser J, Chiu C, Weliky M (2004) Small modulation of ongoing cortical dynamics by sensory input during natural vision. Nature 431:573–578.

Fujisawa S, Buzsáki G (2011) A 4 Hz oscillation adaptively synchronizes prefrontal, VTA, and hippocampal activities. Neuron 72:153–165.

Gao M, Liu C-L, Yang S, Jin G-Z, Bunney BS, Shi W-X (2007) Functional coupling between the prefrontal cortex and dopamine neurons in the ventral tegmental area. J Neurosci 27:5414–5421.

Gao P, de Munck JC, Limpens JHW, Vanderschuren LJMJ, Voorn P (2017) A neuronal activation correlate in striatum and prefrontal cortex of prolonged cocaine intake. Brain Struct Funct 222:3453–3475.

Greenberg DS, Houweling AR, Kerr JND (2008) Population imaging of ongoing neuronal activity in the visual cortex of awake rats. Nat Neurosci 11:749–751.

Hardung S, Epple R, Jäckel Z, Eriksson D, Uran C, Senn V, Gibor L, Yizhar O, Diester I (2017) A functional gradient in the rodent prefrontal cortex supports behavioral inhibition. Curr Biol 27:549–555.

Hunyadi B, Woolrich MW, Quinn AJ, Vidaurre D, De Vos M (2019) A dynamic system of brain networks revealed by fast transient EEG fluctuations and their fMRI correlates. Neuroimage 185:72–82.

Kannathal N, Choo ML, Acharya UR, Sadasivan PK (2005) Entropies for detection of epilepsy in EEG. Comput Methods Programs Biomed 80:187–194.

Karreman M, Moghaddam B (1996) The prefrontal cortex regulates the basal release of dopamine in the limbic striatum: an effect mediated by ventral tegmental area. J Neurochem 66:589–598.

Kenet T, Bibitchkov D, Tsodyks M, Grinvald A, Arieli A (2003) Spontaneously emerging cortical representations of visual attributes. Nature 425:954–956.

Khanna A, Pascual-Leone A, Michel CM, Farzan F (2015) Microstates in resting-state EEG: current status and future directions. Neurosci Biobehav Rev 49:105–113.

Kindler J, Hubl D, Strik WK, Dierks T, Koenig T (2011) Resting-state EEG in schizophrenia: auditory verbal hallucinations are related to shortening of specific microstates. Clin Neurophysiol 122:1179–1182.

Kobak D, Brendel W, Constantinidis C, Feierstein CE, Kepecs A, Mainen ZF, Qi X-L, Romo R, Uchida N, Machens CK (2016) Demixed principal component analysis of neural population data. elife 5.

Laubach M, Amarante LM, Swanson K, White SR (2018) What, if anything, is rodent prefrontal cortex? Eneuro 5.

Lehmann D (1989) Microstates of the brain in EEG and ERP mapping studies. In: Brain Dynamics (Başar E, Bullock TH, eds), pp 72–83 Springer series in brain dynamics. Berlin, Heidelberg: Springer Berlin Heidelberg.

Lehmann D (1990) Brain electric microstates and cognition: the atoms of thought. In: Machinery of the mind (John ER, Harmony T, Prichep LS, Valdés-Sosa M, Valdés-Sosa PA, eds), pp 209–224. Boston, MA: Birkhäuser Boston.

Lehmann D, Faber PL, Galderisi S, Herrmann WM, Kinoshita T, Koukkou M, Mucci A, Pascual-Marqui RD, Saito N, Wackermann J, Winterer G, Koenig T (2005) EEG microstate duration and syntax in acute, medication-naive, first-episode schizophrenia: a multi-center study. Psychiatry Res 138:141–156.

Lehmann D, Ozaki H, Pal I (1987a) EEG alpha map series: brain micro-states by space-oriented adaptive segmentation. Electroencephalogr Clin Neurophysiol 67:271–288.

Lehmann D, Ozaki H, Pal I (1987b) EEG alpha map series: brain micro-states by space-oriented adaptive segmentation. … and clinical neurophysiology.

Liljeholm M, O’Doherty JP (2012) Contributions of the striatum to learning, motivation, and performance: an associative account. Trends Cogn Sci (Regul Ed) 16:467–475.

Luczak A, Barthó P, Harris KD (2009) Spontaneous events outline the realm of possible sensory responses in neocortical populations. Neuron 62:413–425.

Luczak A, McNaughton BL, Harris KD (2015) Packet-based communication in the cortex. Nat Rev Neurosci 16:745–755.

Machens CK, Romo R, Brody CD (2010) Functional, but not anatomical, separation of “what” and “when” in prefrontal cortex. J Neurosci 30:350–360.

Makarov VA, Makarova J, Herreras O (2010) Disentanglement of local field potential sources by independent component analysis. J Comput Neurosci 29:445–457.

Mao BQ, Hamzei-Sichani F, Aronov D, Froemke RC, Yuste R (2001) Dynamics of spontaneous activity in neocortical slices. Neuron 32:883–898.

Massimini M, Huber R, Ferrarelli F, Hill S, Tononi G (2004) The sleep slow oscillation as a traveling wave. J Neurosci 24:6862–6870.

Mathis A, Mamidanna P, Cury KM, Abe T, Murthy VN, Mathis MW, Bethge M (2018) DeepLabCut: markerless pose estimation of user-defined body parts with deep learning. Nat Neurosci 21:1281–1289.

McGinley MJ, David SV, McCormick DA (2015) Cortical membrane potential signature of optimal states for sensory signal detection. Neuron 87:179–192.

Michel CM, Koenig T (2018) EEG microstates as a tool for studying the temporal dynamics of whole-brain neuronal networks: A review. Neuroimage 180:577–593.

Michel CM, Seeck M, Landis T (1999) Spatiotemporal dynamics of human cognition. Physiology (Bethesda) 14:206–214.

Milz P (2016) Keypy –An open source library for EEG microstate analysis. Eur Psychiatry 33:S493.

Milz P, Faber PL, Lehmann D, Koenig T, Kochi K, Pascual-Marqui RD (2016) The functional significance of EEG microstates--Associations with modalities of thinking. Neuroimage 125:643–656.

Milz P, Pascual-Marqui RD, Achermann P, Kochi K, Faber PL (2017) The EEG microstate topography is predominantly determined by intracortical sources in the alpha band. Neuroimage 162:353–361.

Mishra A, Englitz B, Cohen MX (2020) EEG microstates as a continuous phenomenon. Neuroimage 208:116454.

Morales M, Margolis EB (2017) Ventral tegmental area: cellular heterogeneity, connectivity and behaviour. Nat Rev Neurosci 18:73–85.

Muller L, Chavane F, Reynolds J, Sejnowski TJ (2018) Cortical travelling waves: mechanisms and computational principles. Nat Rev Neurosci 19:255–268.

Murray MM, Brunet D, Michel CM (2008) Topographic ERP analyses: a step-by-step tutorial review. Brain Topogr 20:249–264.

Musall S, Kaufman MT, Juavinett AL, Gluf S, Churchland AK (2019) Single-trial neural dynamics are dominated by richly varied movements. Nat Neurosci 22:1677–1686.

Musso F, Brinkmeyer J, Mobascher A, Warbrick T, Winterer G (2010) Spontaneous brain activity and EEG microstates. A novel EEG/fMRI analysis approach to explore resting-state networks. Neuroimage 52:1149–1161.

Okun M, Naim A, Lampl I (2010) The subthreshold relation between cortical local field potential and neuronal firing unveiled by intracellular recordings in awake rats. J Neurosci 30:4440–4448.

Omer DB, Fekete T, Ulchin Y, Hildesheim R, Grinvald A (2019) Dynamic patterns of spontaneous ongoing activity in the visual cortex of anesthetized and awake monkeys are different. Cereb Cortex 29:1291–1304.

Ott T, Jacob SN, Nieder A (2014) Dopamine receptors differentially enhance rule coding in primate prefrontal cortex neurons. Neuron 84:1317–1328.

Palmiter RD (2008) Dopamine signaling in the dorsal striatum is essential for motivated behaviors: lessons from dopamine-deficient mice. Ann N Y Acad Sci 1129:35–46.

Papp EA, Leergaard TB, Calabrese E, Johnson GA, Bjaalie JG (2014) Waxholm Space atlas of the Sprague Dawley rat brain. Neuroimage 97:374–386.

Pascual-Marqui RD, Michel CM, Lehmann D (1995) Segmentation of brain electrical activity into microstates: model estimation and validation. IEEE Trans Biomed Eng 42:658–665.

Passetti F, Chudasama Y, Robbins TW (2002) The frontal cortex of the rat and visual attentional performance: dissociable functions of distinct medial prefrontal subregions. Cereb Cortex 12:1254–1268.

Pasupathy A, Miller EK (2005) Different time courses of learning-related activity in the prefrontal cortex and striatum. Nature 433:873–876.

Peters Y, Barnhardt NE, O’Donnell P (2004) Prefrontal cortical up states are synchronized with ventral tegmental area activity. Synapse 52:143–152.

Petersen CCH, Hahn TTG, Mehta M, Grinvald A, Sakmann B (2003) Interaction of sensory responses with spontaneous depolarization in layer 2/3 barrel cortex. Proc Natl Acad Sci USA 100:13638–13643.

Poulet JFA, Petersen CCH (2008) Internal brain state regulates membrane potential synchrony in barrel cortex of behaving mice. Nature 454:881–885.

Poulsen AT, Pedroni A, Langer N, Hansen LK (2018) Microstate EEGlab toolbox: An introductory guide. BioRxiv.

Raichle ME, MacLeod AM, Snyder AZ, Powers WJ, Gusnard DA, Shulman GL (2001) A default mode of brain function. Proc Natl Acad Sci USA 98:676–682.

Rousseeuw PJ (1987) Silhouettes: A graphical aid to the interpretation and validation of cluster analysis. Journal of Computational and Applied Mathematics 20:53–65.

Sakata S, Harris KD (2009) Laminar structure of spontaneous and sensory-evoked population activity in auditory cortex. Neuron 64:404–418.

Siegle JH, López AC, Patel YA, Abramov K, Ohayon S, Voigts J (2017) Open Ephys: an open-source, plugin-based platform for multichannel electrophysiology. J Neural Eng 14:045003.

Steriade M, Nuñez A, Amzica F (1993) A novel slow (< 1 Hz) oscillation of neocortical neurons in vivo: depolarizing and hyperpolarizing components. J Neurosci 13:3252–3265.

Sul JH, Kim H, Huh N, Lee D, Jung MW (2010) Distinct roles of rodent orbitofrontal and medial prefrontal cortex in decision making. Neuron 66:449–460.

Townsend RG, Gong P (2018) Detection and analysis of spatiotemporal patterns in brain activity. PLoS Comput Biol 14:e1006643.

Uylings HBM, Groenewegen HJ, Kolb B (2003) Do rats have a prefrontal cortex? Behav Brain Res 146:3–17.

Vandecasteele M, M S, Royer S, Belluscio M, Berényi A, Diba K, Fujisawa S, Grosmark A, Mao D, Mizuseki K, Patel J, Stark E, Sullivan D, Watson B, Buzsáki G (2012) Large-scale recording of neurons by movable silicon probes in behaving rodents. J Vis Exp:e3568.

Wackermann J, Lehmann D, Michel CM, Strik WK (1993) Adaptive segmentation of spontaneous EEG map series into spatially defined microstates. Int J Psychophysiol 14:269–283.

Yuan H, Zotev V, Phillips R, Drevets WC, Bodurka J (2012) Spatiotemporal dynamics of the brain at rest--exploring EEG microstates as electrophysiological signatures of BOLD resting state networks. Neuroimage 60:2062–2072.

Zahrt J, Taylor JR, Mathew RG, Arnsten AF (1997) Supranormal stimulation of D1 dopamine receptors in the rodent prefrontal cortex impairs spatial working memory performance. J Neurosci 17:8528–8535.

Zanos TP, Mineault PJ, Nasiotis KT, Guitton D, Pack CC (2015) A sensorimotor role for traveling waves in primate visual cortex. Neuron 85:615–627.

Zhang Y, Pan X, Wang R, Sakagami M (2016) Functional connectivity between prefrontal cortex and striatum estimated by phase locking value. Cogn Neurodyn 10:245–254.

